# *Eubacterium rectale* detoxification mechanism increases resilience of the gut environment

**DOI:** 10.1101/2024.05.09.593360

**Authors:** Tanner G. Richie, Hallie Wiechman, Carson Ingold, Leah Heeren, Abigail Kamke, Sophia Pogranichniy, Kourtney Monk, Trey Summers, Qinghong Ran, Soumyadev Sarkar, Brandon L. Plattner, Ashley Sidebottom, Eugene Chang, Sonny T.M. Lee

**Affiliations:** Division of Biology, Kansas State University, Manhattan, Kansas, United States; Center for Fundamental and Applied Microbiomics, Biodesign Institute, Arizona State University, Tempe, Arizona, United States; Department of Diagnostic Medicine and Pathobiology, Kansas State University, Manhattan, Kansas, United States; Duchossois Family Institute, University of Chicago, Chicago, Illinois, United States; Department of Medicine, University of Chicago, Chicago, Illinois, United States

**Keywords:** Microbiome, gut microbiota, glutathione, microbial derived products, Lachnospiraceae, gut inflammation, shotgun metagenomics, metabolomics, RNA sequencing

## Abstract

Lachnospiraceae members were highly detected in dysbiotic IL-10 KO mice that displayed similar physiological outcomes as control mice. Lachnospiraceae is a highly diverse family of microbes that have been shown to display both commensal and pathogenic characteristics in the colon environment. We investigated the impact of genetic variation in five Lachnospiraceae strains on lowering cellular inflammation and ROS levels. Cell free spent media (CFSM) from *Eubacterium rectale* resulted in lowered ROS, and nitric oxide levels in stressed colon cells. We demonstrated through an array of multi ‘omics and molecular techniques that glutathione (GSH) biosynthesized by *E. rectale* was able to alleviate host ROS damage. We further showed downregulation of cell stress and immune response genes by host RNA sequencing, which is evidence that *E. rectale* microbial products promote recovery and alleviate ROS stress.

**Highlights:**

- Lachnospiraceae detection correlated with lower host inflammation in vertically transmitted dysbiotic mice
- Cell free spent media from *E. rectale* lowered ROS and nitric oxide levels when introduced to stressed Caco-2 cells more effectively than 4 other strains of Lachnospiraceae
- *Eubacterium rectale* spent media products include high levels of glutathione (GSH), carbohydrates, and amino acids
- *Eubacterium rectale* potential gene function for glutathione biosynthesis and metabolism found in the genome
- Lower expression of repair and stress genes of intestinal cells treated with hydrogen peroxide was observed with *E. rectale* CFSM as well as treatment with 15 mM GSH

## Introduction

Microbial diversity (1, 2) is often associated with a healthy gut environment and lower chronic inflammation. A diverse microbiota has the ability to outcrowd potential gut pathogens while stimulating the host immune system, and providing many facilitatory metabolites and nutrients (3). During perturbations, such as antibiotic usage or other medicinal exposures (4), gut infections (5, 6), or autoimmune disorders (7), there is generally a loss in gut microbial diversity, notably from the Firmicutes phylum (2, 8). Firmicutes are a functionally diverse phylum of bacteria, and contain the Lachnospiraceae family, also a diverse and prevalent family at around 10% of the gut microbiota (9). In addition to being a member of the healthy gut microbiota, Lachnospiraceae have also been implicated in gut inflammation alleviation (10, 11) with recent research focusing on butyrate production (12, 13) that is common among members of this bacterial family. The Lachnospiraceae family is a large complex family that has undergone reclassification with several genera being added to this class from Clostridium cluster XIVa (11, 14, 15). Members of this family of bacteria are well known for producing short-chain fatty acids (SCFA) (16), secondary bile acids (17), essential amino acids (18), butyrate (18, 19), and several products involved in reducing radicals and oxidative stress (20).

Many of these Lachnospiraceae derived products are implicated in the alleviation of host gut inflammation by regulating pro-inflammatory cytokine signaling (21), promoting epithelial repair (22), mitigating reactive oxygen species (ROS) damage (23), and maintaining homeostasis (24). However, not all Lachnospiraceae members are metabolically equal, due to considerable metabolic and functional diversity within the species (14) resulting in wide variance of metabolism in each species of the Lachnospiraceae family (25). Some species do not produce butyrate (11), and others have a variety of virulence genes (26) including antimicrobial resistance (27) and defensive genes (28). Moreover, the presence of some species may reduce gut inflammation in disease states like ulcerative colitis (UC) but not in inflammatory bowel disease (IBD) (29). This suggests that a specific community of Lachnospiraceae producing a wide range of beneficial metabolites and products are critical in lowering inflammation and promoting gut repair (25, 30). The lack of a clear understanding of how Lachnospiraceae derived metabolites could alleviate gut inflammation hinders the ability to develop targeted therapeutic interventions and diagnostic tools. In addition, it raises further questions about the specific mechanisms by which these metabolites interact with the host immune system to reduce inflammation. This study explored the microbial functions of Lachnospiraceae to identify possible metabolites that contribute to lowering inflammation or ROS damage.

Host-derived glutathione (GSH) is well established in its antioxidant capability within the healthy gut, and its function as a node in the circuitry of intestinal signaling with connections to host-cell proliferation and differentiation (31, 32). During chronic gut inflammation, host GSH synthesis is often impaired, leading to worsened outcomes (33). Thus, GSH serves as a critical biomarker for host health and is important for promoting homeostasis and gut repair. This study builds upon existing theories that certain bacterial populations within the Lachnospiraceae family are able to induce host GSH synthesis through the production of SCFA, and further showed that Lachnospiraceae can utilize cysteine, glutamic acid, and glycine, which are needed to biosynthesize GSH (34) to aid in host recovery. Microbial derived GSH could aid in alleviating ROS and facilitates host-gut epithelial repair when host-derived GSH is impaired due to damage and immune response. Thus, the involvement of Lachnospiraceae-derived amino acids, used in biosynthesis of GSH, that promotes gut homeostasis through eliminating ROS are of essential importance for designing probiotics or other therapeutics (34, 35).

Here, we conducted an in-depth characterization of Lachnospiraceae derived metabolites, and their ability to alleviate gut inflammation and ROS damage, which provides insights into the immune response of the inflamed host after exposure to microbial metabolites. This study combines a multi-omic *in vivo* approach with microbiological and molecular *in vitro* techniques to propose metabolic pathways of interest in Lachnospiraceae that could aid in alleviation of the inflamed gut. We observed that the Lachnospiraceae derived metabolites have unique metabolic profiles that support alleviation of inflammation; however, metabolites derived from *Eubacterium rectale* lowered cellular stress more effectively than the other strains. We observed a significant reduction of GSH in the cell-free spent media of *E. rectale* compared with the other strains. Following that, we focused on the metabolic mechanisms of *E. rectale* derived GSH, and incorporated the host response to shed light on host-microbe interactions and mechanisms that may be involved with stress relief. Our findings mechanistically link associations with Lachnospiraceae derived GSH, other crucial microbial metabolic pathways, and host’s immune markers of gut health to provide insights into *E. rectale*’s role in potential protection against gastrointestinal disorder.

## Results

Fecal content from pups at weeks 4, 6, 9, and 24 of age, were used for shotgun metagenomic sequencing, to capture development and effects of inherited dysbiosis as well as before and after treatment with dextran sulfate sodium (DSS). Mice in the dysbiotic group displayed vertically transmitted dysbiosis from dam exposure to cefoperazone, and murine pups developed for 23 weeks until 2.5% DSS was administered at the endpoint of the study to induce colitis in all mice. Control mice originated from untreated IL-10 knockout mice and were monitored for 23 weeks until DSS exposure in drinking water (Supplementary Table S1). We used genome-resolved metagenomics for high resolution characterization of the gut microbiota, combined with host gene expression and microbial metabolite analyses to explore the impact of Lachnospiraceae members on host recovery from dysbiosis.

Shotgun metagenomic sequencing of fecal samples resulted in over 1,000,437,831 sequences with an average of approximately 14,000,000 reads per sample. Assembly of sequences from the dysbiotic group (n = 30) and control group (n = 45) resulted in 439,024 contigs longer than 1,000 nucleotides. We observed a clear pattern of vertically transmitted dysbiosis belonging to several microbial classes across each sample compared to the control mice immediately after cefoperazone exposure ended (Supplementary Figure S1). Our ultimate goal of this study was to monitor the outcomes of vertically transmitted dysbiosis throughout mouse development, and investigate potential microbial members and their mechanisms influencing recovery.

### Dysbiotic mice showed minimal differences from control mice in colon histology and cytokine markers

To gain insights into the host response after vertically transmitted dysbiosis, we performed histology and a broad cytokine panel on murine colons and serum respectively (Supplementary Table S1). As seen in previous studies, we expected dysbiotic mice with induced colitis to display more severe clinical signs of inflammation compared with control mice with induced colitis, which highlights the importance of a diverse gut community on immune regulation (36–38). Surprisingly, we observed that offspring from dysbiotic dams and control mice displayed similar low grade colon inflammation with signs of neutrophil infiltration and slight surface damage at week 24, due to the chemically induced colitis from DSS (Figure 1A). Overall, dysbiotic mice were not significantly different from mice in the control group in all grading categories including inflammation, surface damage, and crypt hyperplasia scores (Figure 1B). It was surprisingly evident that the dysbiotic mice might have recovered from dysbiosis during development as there are no significant differences in colon inflammation between groups.

**Figure 1.**
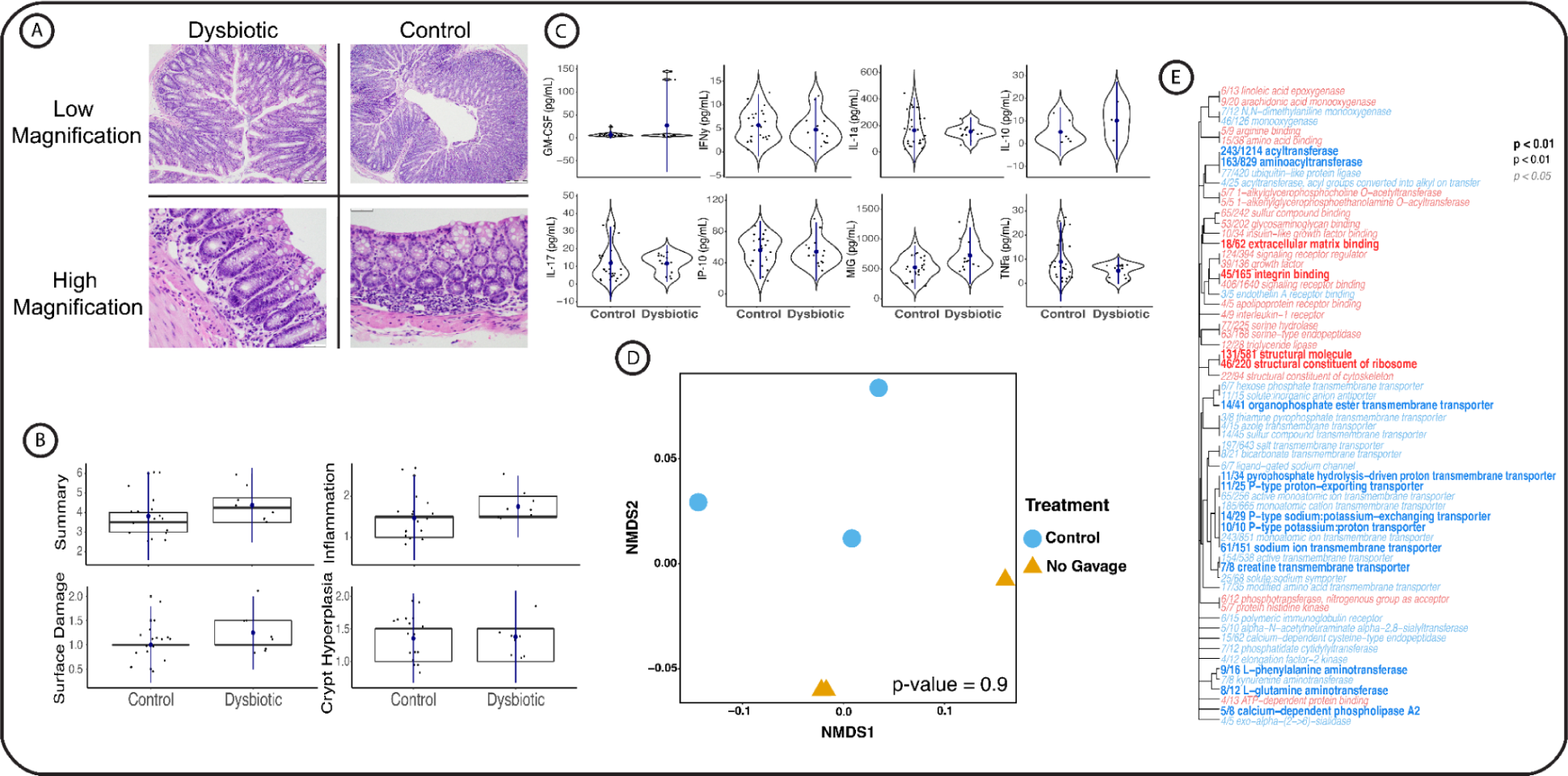
Dysbiotic mice display similar histology scores, metabolite profiles, and gene expressions as control mice. (A) Histology images at low magnification and high magnification in both groups of mice. (B) Boxplots of histology scores for Summary (0-9) scores and broken into categories of Inflammation (0-3), Surface Damage (0-3) and Crypt Hyperplasia (0-3) with no statistical differences between treatment groups. (C) Murine serum cytokine violin plots of cytokines typically highly expressed during colon inflammation also result in no statistical difference between groups. (D) NMDS plot of mouse metabolite profiles resulting from PERMANOVA indicate no metabolic differences between treatments. (E) Gene Ontology analysis of gene family expressions in both treatment groups. Statistical difference marked by bolding and text size with blue colored text corresponding to downregulation of a gene family in dysbiotic mice, red text is upregulation of a gene family in dysbiotic mice compared to control. Few gene classes are significantly different with a p-value of < 0.05 between treatments with no clear immunological patterns observed. Indicating the dysbiotic mice by the end of the study period display colon inflammation levels similar to control mice.

We further evaluated serum systemic markers using a 31 multiplex pro-inflammatory cytokine panel for all mice that survived until week 24. Similar to our observations regarding colon inflammation, 30 of the 31 cytokines measured displayed no significant difference between dysbiotic and control mice (Figure 1C). Monokine induced by gamma interferon (MIG) was significantly higher in dysbiotic mice (p = 0.0061) than in control mice. MIG (also referred to as CXCL9) is a chemokine that is implicated in gut inflammation, however is usually accompanied with other cytokines including IL-1ɑ, IL-10, IL-17, and TNF-ɑ during an immune response (39). MIG is also activated in response to the introduction of a gut microbiota in both murine and cell models (40). Because this was the only cytokine with elevated levels, we suspect this increased level of MIG in dysbiotic mice was due to the reappearance of gut microbes during development (41, 42). In summary, we see little differences in cytokine profiles between treatment groups. Thus, we surmise if there was a microbial associated mechanism involved in the resulting similarity in cytokine profiles of the dysbiotic mice and the control mice in our study.

To ensure a persuasive hypothesis that the dysbiotic mice displayed a similar host status to control mice, we conducted a global metabolite panel and host RNAseq from each treatment group. We used PERMANOVA analysis and showed that there were no significant differences among the metabolite profiles between treatment groups (p-value = 0.9 and error margin = 7.1 x 10^-5^, Figure 1D). To gain perspective on the host response, RNA sequencing on mouse colons from each treatment also resulted in minimal gene family regulation changes, with 63 significantly regulated gene families between treatments (Figure 1E). These data suggest there is not a clear difference of gene family regulation, and together with the lack of inflammatory gene upregulation, point to the recovery of the mice in the dysbiotic group.

### Increase in Lachnospiraceae bacterial populations in dysbiotic mice associated with lowered host pro-inflammatory cytokine

The following work stems from the fact that there were no statistical differences in the histology, cytokine expressions, metabolite profiles and host gene regulations, which suggests mice recovered from vertically transmitted dysbiosis. We were then curious if the gut microbiota played a part in the similar status of these dysbiotic mice to control mice. Therefore, to characterize the gut microbiota of dysbiotic mice, we performed shotgun metagenomic sequencing on the fecal samples at weeks 4, 6, 9, and 24 of age (Supplementary Table S2). We resolved 158 non-redundant metagenomes-assembled genomes (MAGs) (Supplementary Figure S1), with 73 MAGs annotated to the Lachnospiraceae family (Supplementary Table S2). We noticed Lachnospiraceae MAGs displayed an interesting longitudinal pattern in the dysbiotic mice (Figure 2A). Although low in detection until week 6 (Supplementary Table S2; Figure 2A and 2B), we saw a significant increase in Lachnospiraceae in the dysbiotic mice after week 9 (Supplementary Table S2; Figure 2A and 2B). Control mice, on the other hand, did not display this pattern (Supplementary Table S2; Figure 2A). Interestingly, we noticed that control mice had the highest detection of Lachnospiraceae at week 6, which implicated their presence earlier in life as opposed to the dysbiotic group where this increased detection was seen later at week 9 of age.

**Figure 2.**
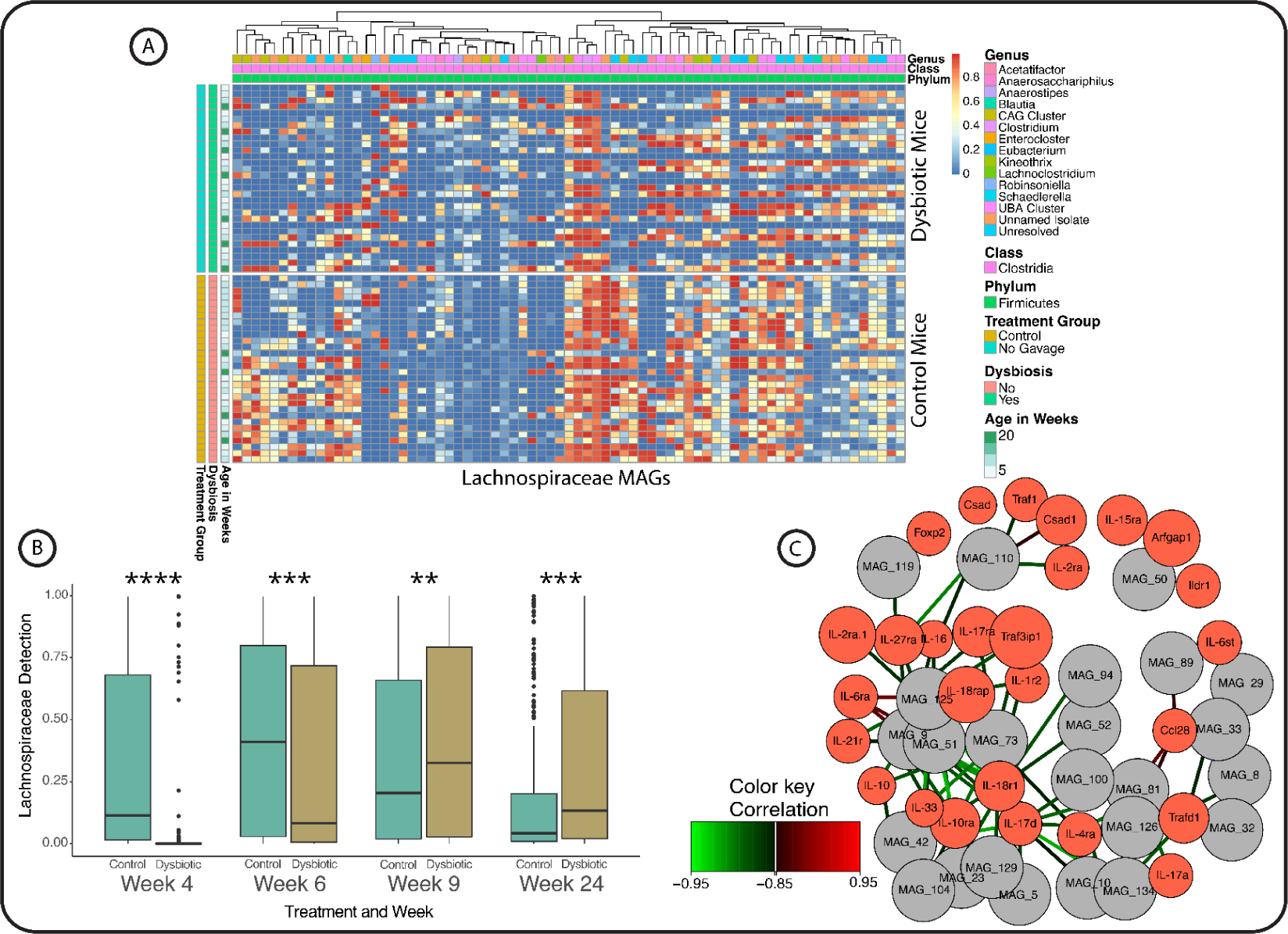
Increased Lachnospiraceae detection over murine development correlated with downregulation of pro-inflammatory genes. (A) Clustered heatmap with rows corresponding to mouse samples sorted by treatment group and age in weeks. Constant detection levels are observed in control mice, however over time, Lachnospiraceae increase over time in the dysbiotic mice. (B) Boxplot of Lachnospiraceae MAG detections by week comparing control mice (Green) to Dysbiotic mice (Gold). Significant p-values marked by asterisks * < 0.05, ** < 0.005, *** < 0.0005, and **** < 0.00005. (C) Network analysis of discriminant analysis of RNA sequencing gene counts (Red) coupled with MAG detection levels (Gray). A correlation cutoff of 0.95 was used to only visualize strong associations with (Green) lines indicating a negative correlation, and (Red) lines marking a positive correlation to the detection of a MAG with the upregulation of a gene.

We wanted to determine if an increase in Lachnospiraceae populations were associated with beneficial host response patterns. Thus, to link this pattern with the recovery of the dysbiotic mice, we performed a discriminant analysis on MAG detections and host RNA sequencing counts. Gene counts for pro-inflammatory cytokine receptors, including IL-17, IL-10, IL-6, and IL-1 that are often highly expressed in inflammatory bowel disease (43), were negatively correlated with the higher detection of Lachnospiraceae MAGs (Figure 2C). Our results indicated that Lachnospiraceae MAGs might interact with the host immune system to lower pro-inflammatory cytokine expression, which effectively reduced the host’s immune responses (44, 45). However, with the high diversity and heterogeneity within the Lachnospiraceae family, it is difficult to dissect which members of Lachnospiraceae are driving the alleviation of inflammation. In the next sections, we examined all 73 murine MAGs, and compared them to five Lachnospiraceae strains that are implicated with a healthy gut, to further drive at a mechanism Lachnospiraceae employ in promoting gut homeostasis.

### Metabolic pathways of Lachnospiraceae MAGs included antioxidant and secondary metabolite productions that may benefit host

With the Lachnospiraceae bacterial populations displaying a unique detection pattern longitudinally in our study, we further investigated the potential functions of these 73 MAGs that could result in possible host recovery. We used Bacterial and Viral Bioinformatics Resource Center (BV-BRC) genome annotation and subsystems to visualize and quantify the metabolic potential of all 73 MAGs. The highest number of genes by pathway for all MAGs was GSH metabolism with 225 genes found among the 73 MAGs, and resulted in 100% coverage of the pathway (Figure 3A). Six other pathways included in the analysis involved the metabolism of amino acids, including glycine, serine, threonine, arginine, proline, lysine, and cysteine. Amino acid metabolism within microbes is often the beginning step toward production of SCFA with the amino acids mentioned above being involved in the production of butyrate (46, 47). With GSH metabolism being the highest represented pathway, we next investigated how microbial GSH assimilation could impact host outcomes. The following section will discuss the rationale for choosing Lachnospiraceae isolates that are relevant to human health and have similar metabolic products to the murine derived MAGs.

**Figure 3.**
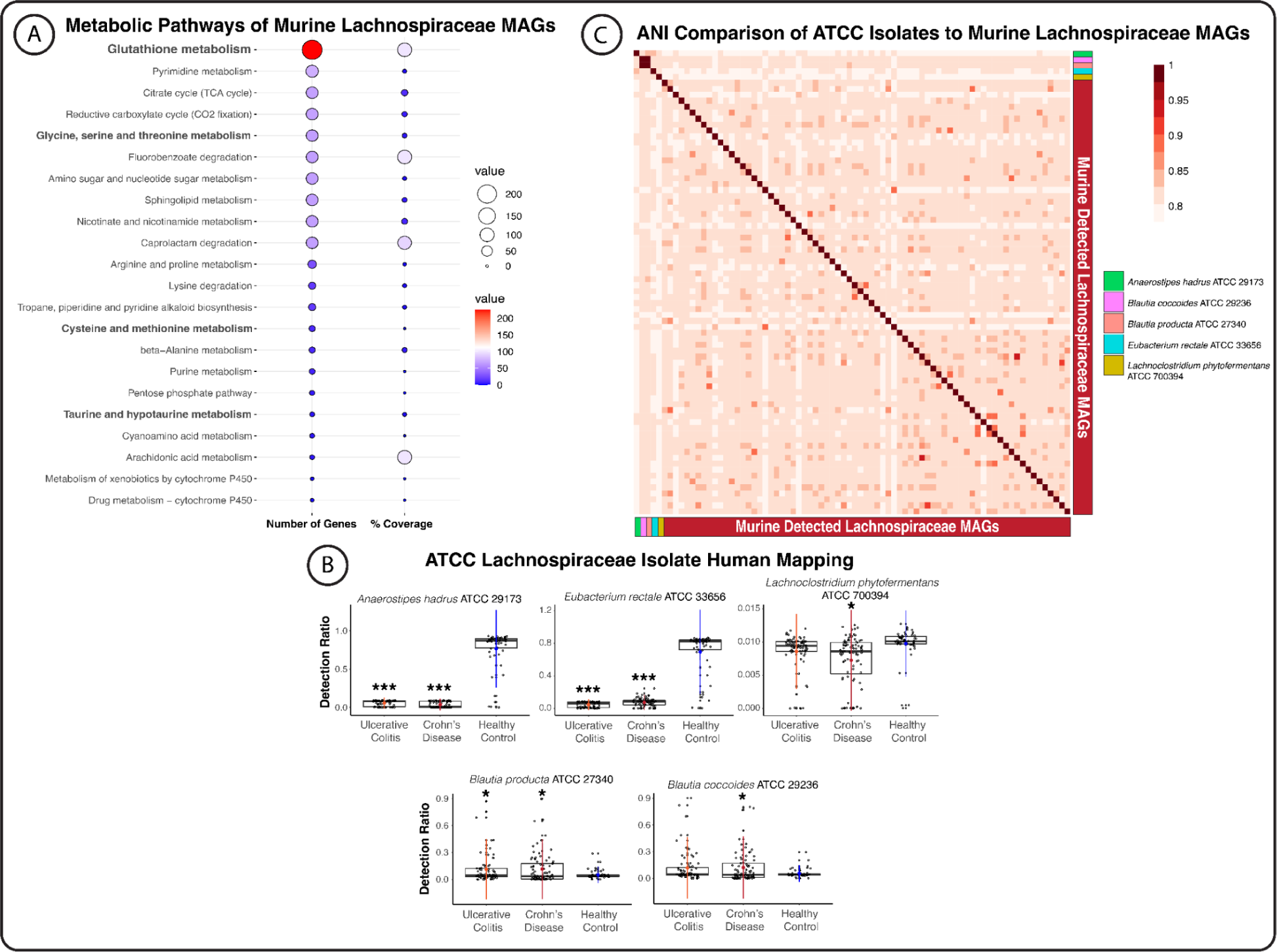
Genus level match of Lachnospiraceae MAGs to available isolates, presence of several beneficial pathways to the host, and isolates are present in healthy patient samples and much lower in diseased patients. (A) Balloon plot of metabolic pathway summary of all 73 murine Lachnospiraceae MAGs. Size of the balloon indicates either the number of genes in the corresponding pathway or the percent coverage of the entire pathway shown. (B) Mapping the genomes of the available 5 ATCC isolates to human samples from a subset of the Discovery cohort indicated patients with Ulcerative colitis and Crohn’s Disease were significantly lower in almost every strain. Significant p-values were marked with asterisks with * < 0.05, ** < 0.005, *** < 0.0005. (C) Average Nucleotide Identity (ANI) resulted in around an 80-90% match of available Lachnospiraceae to murine detected MAGs. Murine detected isolates are indicated in Red, the available isolates are marked in the legend.

### Lachnospiraceae members are detected higher in healthy human samples as an indicator of beneficial members

Most MAGs resolving to the Lachnospiraceae family were not annotated to species level, except for the genera *Blautia*, *Acetatifactor*, *Anaerostipes*, *Clostridium*, *Eubacterium*, *Enterocloster*, *Robinsoniella*, *Kineothrix*, *Lachnoclostridium*, *Anaerosacchariphilus*, and *Schaedlerella* (Supplementary Table S2). An average nucleotide identity (ANI) (Supplementary Table S3) resulted in Lachnospiraceae MAGs matching around 80-90% to the available ATCC isolates, indicative of matching at the genus level but not reaching 95% threshold for likely species level identification (48) (Figure 3B). With such wide variety among the isolates and the MAGs, we obtained all five strains from ATCC that were compared to the MAGs seen in dysbiotic mice; *Anaerostipes hadrus* B2-52, *Blautia coccoides* CLC-1, *Blautia producta* VPI 4299, *Eubacterium rectale* VPU 0990, and *Lachnoclostridium phytofermentans* ISDg. With isolates spanning 3 genera and a set within the same genus, these isolates provide insights into functional differences and determine which strain has more potential to alleviate host inflammation.

In order to understand the prevalence of these strains in human populations, we mapped the isolate’s genomes to the PRISM cohort study of IBD patients diagnosed with ulcerative colitis (UC) and Crohn’s disease (CD) (Supplementary Table S3), compared to non-IBD patient samples (49) (Figure 3C). *Anaerostipes hadrus* was detected significantly higher in the healthy control samples (p-value < 0.05), but not in the UC group and the CD group (p-value = 0.57). *Lachnoclostridium phytofermentans* was detected significantly higher in the healthy control samples as compared to the CD group (p-value = 0.00001), but not statistically significant when compared to the UC group (p-value = 0.089). Interestingly, the *Blautia* genus displayed a different detection pattern. *Blautia coccoides* was detected at lower ratios in the non-IBD control samples; however, only compared to the CD group (p-value = 0.032), not the UC group (p-value = 0.059). *Blautia producta* was also detected lower in the non-IBD control samples compared to the UC group (p-value = 0.043) and in the CD group (p-value = 0.027). *Eubacterium rectale* was detected much higher in the non-IBD control samples than the UC group (p-value < 0.05) and the CD group (p-value < 0.05). A higher prevalence in the healthy group versus both disease groups suggests that these microbes may be involved in maintaining homeostasis, and may also play a role in recovery from inflammation. These results clearly indicate the presence of Lachnospiraceae populations in control samples, and are depleted in patients with inflammatory diseases, which suggests the importance for the presence of these strains of Lachnospiraceae, especially *A. hadrus* and *E. rectale*.

### Species level differences in gut inflammation alleviation in vitro using induced inflammation and ROS damage

With the understanding that each Lachnospiraceae strain is functionally diverse across multiple metabolic pathways, we investigated how these differences could affect host inflammatory outcomes. Lachnospiraceae have long been known as major producers for SCFA and butyrate (13, 50), which can help in gut repair and lowering oxygen levels in the colon and establishing local microenvironment for inflammation recovery. We surmised that each of the five Lachnospiraceae strains in this study has unique metabolites that may impact the ability to aid in host recovery from inflammation resulting from two separate exposure tests to DSS or hydrogen peroxide (H_2_O_2_). These two inflammatory exposures either DSS or H_2_O_2_ were utilized because the immune mechanisms affected by DSS are strikingly similar to colitis (51, 52), whereas H_2_O_2_ causes ROS damage often inflicted by more aerobic pathogenic gut microbes present in the dysbiotic gut (53, 54). Because GSH is an antioxidant (35), we also sought to determine if GSH alone is capable of reducing cellular inflammation or peroxide stress to the physiological level we observed in our study.

Inflammation and cell death were quantified by measuring nitric oxide (55, 56), an immune system product indicative of an immune response, quantified ROS levels, which indicates host damage and immune response, and cytotoxicity which captured cell death (57, 58) (Supplementary Table S4). We observed that *E. rectale* (p-value = 1.4 x 10^-6^) and *B. coccoides* (p-value = 4 x 10^-7^) cell-free spent media (CFSM) significantly lowered nitric oxide levels compared to DSS inflamed cells, while none of the strains lowered nitric oxide levels in the H_2_O_2_ exposed cells (Figure 4A). We further noticed that GSH did not significantly lower nitric oxide levels at any concentration in either stress exposure (Figure 4B). Our result suggests that GSH may not contribute to lowering the immune response; however, it is likely involved with alleviating ROS damage. On the other hand, CFSM from *E. rectale* and *B. coccoides* may be involved in diminishing the immune response.

**Figure 4.**
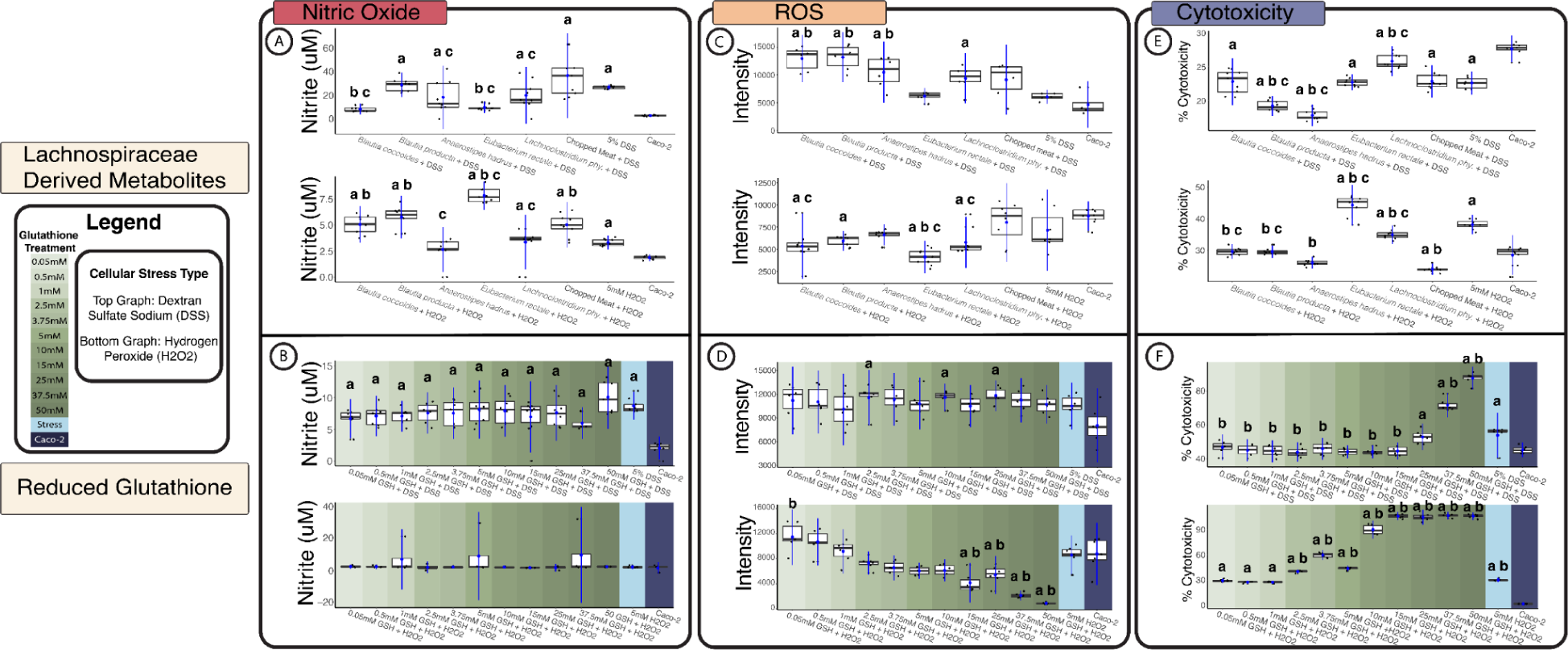
Molecular assays quantifying immune status, ROS damage, and cytotoxicity of each Lachnospiraceae strain’s spent media or reduced glutathione (GSH) on Caco-2 cells inflamed by 5% DSS or ROS stressed with 5 mM hydrogen peroxide (H2O2). (A) Griess assay was used to quantify the immune response by-product nitric oxide. Boxplots show *B. coccoides* and *E. rectale* are significantly lower than the inflamed cell control and chopped meat inflamed control for DSS, and no strains significantly lowered nitric oxide levels when cells are stressed in H2O2. (B) GSH was tested at several half log concentrations in cells stressed with DSS and H2O2 with no statistical significance in lowering nitric oxide in either stressor. (C) Boxplots showing ROS levels, highlight no strain’s spent media lowered ROS levels in DSS inflammation, however *E. rectale*, *B. coccoides*, and *B. producta* were significantly lower than the chopped meat inflamed control in H2O2 stress. (D) GSH at any concentration did not lower ROS levels in DSS stress, but at 15 mM GSH and higher concentrations was significantly lower than H2O2 treated cells. (E) Cytotoxicity boxplots highlight the spent media lowered toxicity in DSS inflamed cells, with *A. hadrus*, and *B. producta* treated cells having significantly lower cytotoxicity than inflamed control. However with H2O2 treated cells, cytotoxicity *B. coccoides*, *B. producta*, and *A. hadrus*, and *E. rectale* displayed higher cytotoxicity than the H2O2 treated control. (F) Increased levels of GSH over 15 mM induced high levels of cytotoxicity in both treatments, with lower levels showing no difference from control cells. Significance (p-value < 0.05) symbolized by comparison (a = significantly different from Caco-2 cells only), (b = significantly different from the stress control DSS or H2O2), and (c = significantly different from Chopped meat media control with stressor DSS or H2O2). Figure legend on the left side of the graph indicates the color scale representing the concentration of GSH, and indicates the top graph is testing DSS and the bottom graph in each subfigure tested H2O2.

CFSM from *E. rectale* displayed the lowest ROS levels with DSS stress, and was not significantly different from control cells while other strains (*B. coccoides*: p-value = 0.0002; *B. producta*: p-value = 0.0001; *A. hadrus*: p-value = 0.038) displayed significantly higher ROS levels in the DSS exposed cells. In H_2_O_2_ exposed cells, *E. rectale* (p-value = 1.8 x 10^-6^), *L. phytofermentans* (p-value = 0.041), and *B. coccoides* (p-value = 0.0074) showed significantly lower ROS levels than the H_2_O_2_ control (Figure 4C). While there are studies on microbial produced GSH (59, 60), our results illuminate the genetic differences among the various members in the Lachnospiraceae family, in the biosynthesis of GSH, with *E. rectale* shown as the largest producer of GSH. There were no significant differences in ROS levels stressed with DSS for all concentrations of GSH; however, concentrations at and higher than 15 mM (p-value = 9 x 10^-5^) did lower ROS significantly with H_2_O_2_ stressed cells (Figure 4D). This suggests that a high amount of GSH (≥ 3.75 mM) is involved in lowering ROS levels when a ROS stress is implemented, but adding GSH did not lower ROS levels when there was an inflammation stressor. Cytotoxicity remained low for all DSS exposed cells; however, *B. producta* (p-value = 3.8 x 10^-7^) and *A. hadrus* (p-value < 0.05) CFSM were significantly lower than the inflamed control. H_2_O_2_ exposed cells revealed similarly that *B. coccoides* (p-value < 0.05), *B. producta* (p-value < 0.05), and *A. hadrus* (p-value < 0.05) were all significantly lower than the stressed control, with *E. rectale* (p-value < 0.05) CFSM significantly higher in cytotoxicity (Figure 4E). We also noticed that GSH showed an interesting pattern of increasing cytotoxicity after 25 mM for DSS stressed cells, and over 10 mM in the H_2_O_2_ exposed cells (Figure 4F). This is likely due to the low pH shift in solution after 15 mM GSH, while most cellular concentrations of GSH range from 1-10 mM (32). In summary, our results highlight that in both stressors and outputs of inflammation, *E. rectale* CFSM was more effective in lowering levels of both nitric oxide and ROS, whereas other strains, like *B. producta* and *A. hadrus,* might have more protection against cytotoxicity rather than lowering inflammation and ROS damage. Here we also showed that GSH is able to lower ROS levels significantly, but does not significantly impact nitric oxide levels, which further stresses its importance as a ROS alleviator and antioxidant at cellular concentrations.

### Functionally diverse Lachnospiraceae have inflammatory alleviating metabolites and antioxidant GSH

With the results showing the CFSM of Lachnospiraceae alleviated cell inflammation, next we investigated the microbial products of these five isolates. We used a widely targeted metabolite analysis to investigate the metabolic profile of CFSM from Lachnospiraceae, and to identify potential metabolic pathways that contribute to the alleviation of host inflammation (Supplementary Table S5). We observed that amino acids had the highest represented class at 56% of all detections in every Lachnospiraceae metabolite profile, with other homeostasis promoting classes (61) including fatty acids (3.5%), bile acids (1.2%), and carbohydrates (4.4%) also represented. Next, a comparison of metabolites of each Lachnospiraceae strain CFSM revealed a core of 82 metabolites in common. However, each strain consisted of unique metabolic products *Blautia coccoides* (n = 6), *Blautia producta* (n = 7), *Lachnoclostridium phytofermentans* (n = 6), *Anaerostipes hadrus* (n = 15), and *Eubacterium rectale* had the highest number of 47 unique metabolites compared to the other strains (Figure 5A). *Eubacterium rectale* specific metabolites include L-tryptophan, L-glutamine, L-histidine, and isobutyric acid. Two of the 47 unique metabolites also include oxaloacetic acid and indole-3-acetic, which are responsible for fatty acid synthesis and oxidative stress reduction (62), and known to alleviate colitis associated immune responses, respectively (63, 64). The distinct metabolic profile of *E. rectale* CFSM provides some evidence that this strain may contribute to decrease inflammation by production from multiple metabolic pathways attributed to lower host inflammation.

**Figure 5.**
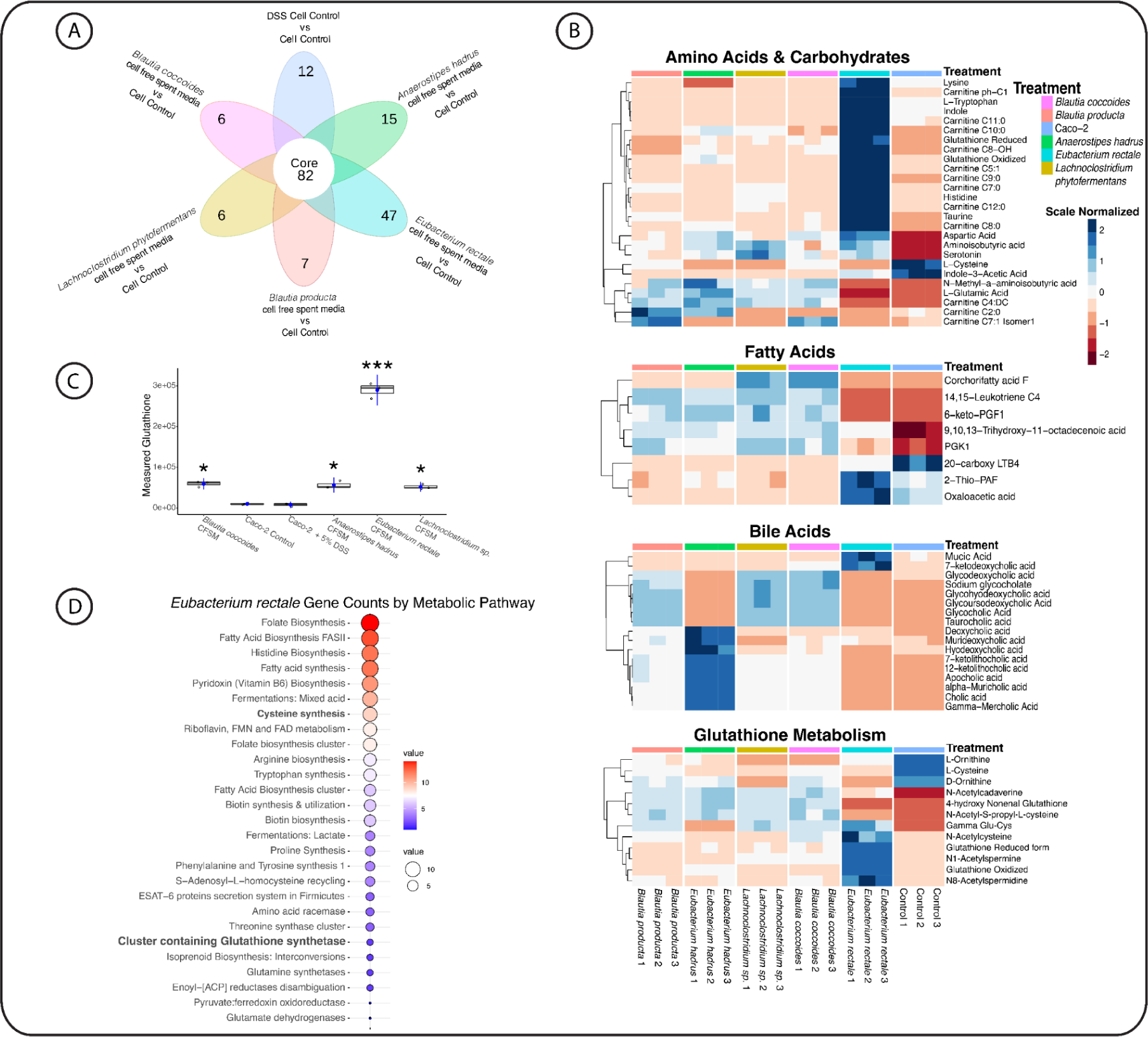
Metabolomic analysis of five Lachnospiraceae strains spent media indicative of functional diversity in microbial family, large production of amino acids, SCFA and bile acids. (A) Venn diagram of all five strains and a Caco-2 cell control, showcasing the unique metabolites of each strain’s spent media, with *Eubacterium rectale* having 47 unique metabolites from the other four strains. (B) Clustered heatmaps displaying relevant metabolites in gut repair including amino acids, glutathione pathways, bile acids and short chain fatty acids in all five strains. Normalized scale bars to the right of each graph indicate the observations of each sample. Notably, *E. rectale* spent media consists of a variety of metabolites as well as having some of the highest counts of metabolites including glutathione, indole, taurine, and several carnitine derivatives. (C) Balloon map of gene subsystems counts for E. rectale by metabolism pathway highlighting potential functional capacity of production of GSH as well as several amino acids with circle size and color indicating number of genes in the pathway. (D) Boxplot of GSH levels in each sample from metabolomic analysis with significance marked by a p-value of * < 0.05, ** < 0.005, *** < 0.0005.

We investigated the metabolic profiles of each Lachnospiraceae strain to determine which strains are major producers of metabolic classes related to host recovery. *Eubacterium rectale* was a major producer with the highest counts of carnitine isomers, histidine, L-lysine, glutathione, and L-taurine compared to the four strains (Figure 5B). Fatty acids were produced highly by all five strains of Lachnospiraceae with *E. rectale* producing the highest amount of 2-thio-PAF and oxaloacetate acid. Bile acids were also produced by all five strains, with *A. hadrus* being the major producer, and *E. rectale* having high measurements for mucic acid and 7-ketodeoxycholic acid. Finally we examined GSH pathway metabolites, and showed that all five strains have partial production of some key metabolites, like N-acetyl-S-propyl-L-cysteine, D-ornithine, and nonenal glutathione. However, *E. rectale* produced the endpoint metabolites including N1-acetylspermidine, N-acetylcysteine, and both reduced and oxidized forms of glutathione (Figure 5B).

We used metabolomics and cellular assays to show that *E. rectale* have strong impacts on cell health, and produces several metabolites known for their antioxidant and gut repair properties, including L-carnosine, L-cysteine, glutathione, L-histidine, and L-taurine. L-carnosine is involved with gut integrity and wound repair (65), while L-cysteine is implicated in gut lining repair (66, 67). L-histidine is implicated in suppression of pro-inflammatory cytokines and plays a role in diminishing colitis (68) in addition to L-taurine, which promotes goblet cell formation and aids in tight junction regulation (69, 70). L-cysteine, L-histidine, and L-taurine have also been considered for supplementation to patients with IBD as these patients often have very low serum levels of these amino acids (33, 71). GSH is a well-documented antioxidant aiding in ROS damage and driver of an anaerobic gut environment (31, 32). We quantified the GSH levels across strains and the colon cell controls, showing there was significantly more GSH in *E. rectale* than the other four strains of Lachnospiraceae (Figure 5C). Our results strongly suggest that the highly effective ROS alleviating characteristics of *E. rectale* was in part due to the high production level of GSH. We further confirmed the biosynthesis capacity of GSH by *E. rectale*, and found several biosynthesis pathways of GSH, as well as the precursor amino acids like L-cysteine synthases (Figure 5D). We also noticed several other microbial metabolism mechanisms that can aid to alleviate host inflammation, such as fatty acid synthesis, tryptophan, and several other amino acid biosynthesis pathways. We next focused on *E. rectale* and its impact on host response, as well as investigation of *E. rectale* derived GSH on alleviation of cellular inflammation. To provide insights into the rescue effect *E. rectale* has on host inflammation.

### Cell response of peroxide damaged Caco-2 cells exposed to Eubacterium rectale CFSM and reduced glutathione showed downregulation of defense and repair genes

Expanding on the cellular assay experiment, we investigated the global host response when exposed to H_2_O_2_, compared to the response of ROS stress with treatment of *E. rectale* CFSM. We evaluated the antioxidant GSH in lowering ROS cell stress because it is a microbial product from *E. rectale,* and is protective against ROS damage. We observed patterns of highly differentially regulated genes in each treatment comparison (Supplementary Table S6). We performed a gene ontology analysis (GO) to investigate gene class differential regulation, and showed that only four gene families, glycosyltransferase, hydrolase, RNA binding, and carbohydrase kinase, were (n = 3) upregulated in the *E. rectale* CFSM treatment. This demonstrated a differential regulation between ROS stressed cells treated with *E. rectale* CFSM compared to control, unstressed cells (Figure 6A). Our results indicate that treating ROS stressed cells with *E. rectale* CFSM significantly reduced host damage, and upregulated gene repair pathways in ROS-stressed cells. Compared with *E. rectale* CFSM treatment, ROS-stressed cells treated with 15 mM of reduced GSH revealed 20 gene classes differentially regulated, with classes including transferases, signal receptor binding, electron transfer, and extracellular matrix structure constituents. Comparing the H_2_O_2_ treated with GSH and the unstressed controls, 18 classes were differentially regulated (Figure 6A). GSH regulation patterns were different compared with the *E. rectale* CFSM; however, they also displayed a differential gene class pattern from inflamed cells (Supplementary Figure S2). GSH treated cells exposed to H_2_O_2_ indicated 31 gene classes differentially upregulated (n = 17) compared to inflamed controls. Our data suggest that GSH increased the ability of different gene family pathways to reduce ROS damage to host cells, compared to *E. rectale* CFSM.

**Figure 6.**
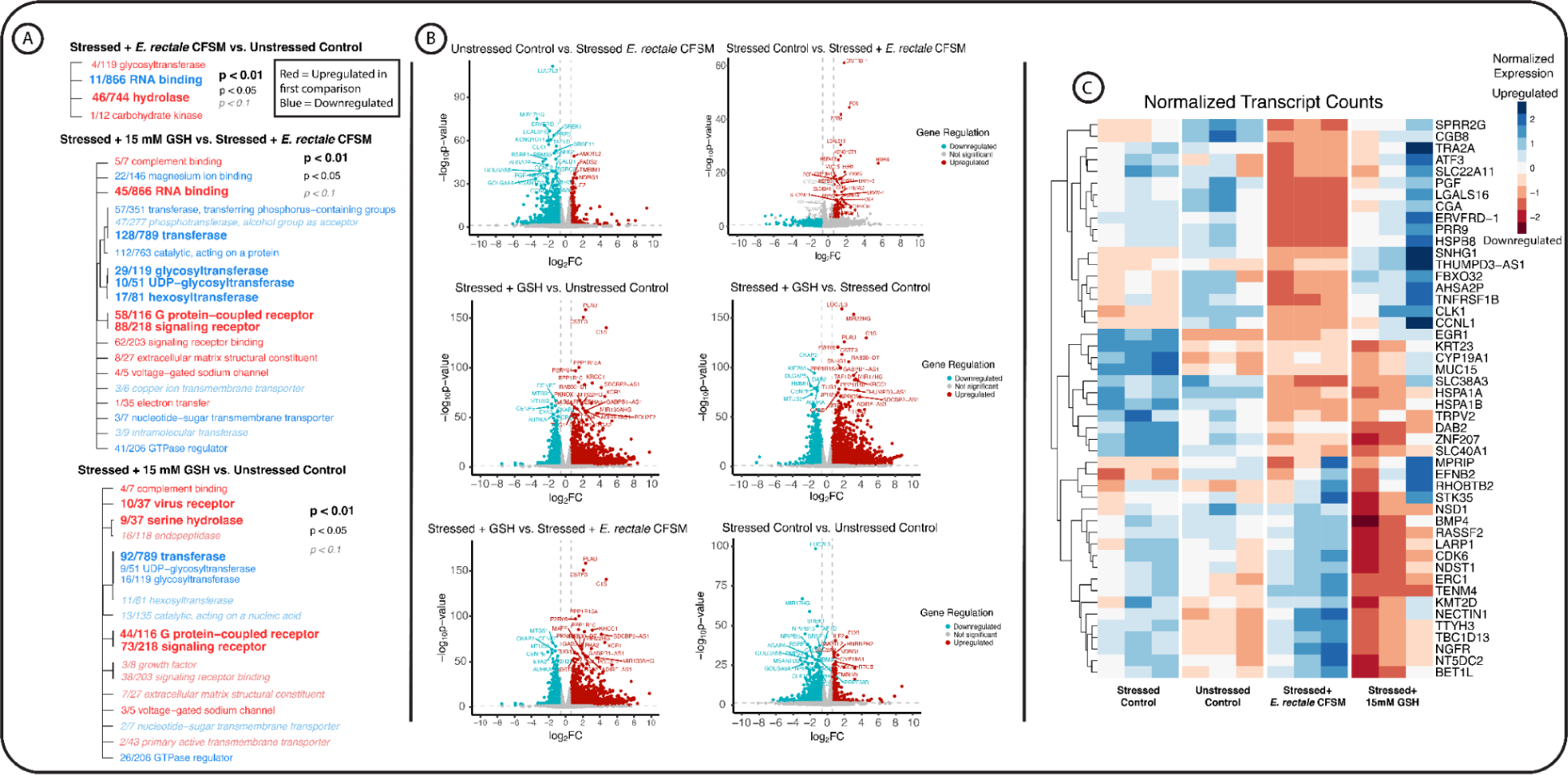
RNA sequencing of peroxide stressed Caco-2 cells treated with CFSM of *E. rectale* resulted in little gene differentiation from uninflamed Caco-2 cells, and treatment with reduced GSH reduced several damage repair genes and environmental sensing genes. (A) Volcano plots of DEseq2 comparison analyses with a p-value < 0.05 and fold change of > 0.6 with the top 30 differentially expressed genes labeled on each plot. Color scale depicts non-significantly regulated genes as gray, upregulated genes as red and downregulated genes as blue. (B) Gene ontology analysis between treatment group comparisons indicating gene families that are differentially regulated based on p-values less than 0.05 and indicated with text size and fonts. (C) Clustered and normalized heatmap by row of gene count data. The top 50 significantly different genes between stressed cells and *E. rectale* CFSM versus 15mM GSH.

We identified differential host gene isoforms analysis to provide a higher resolution of the host response. Volcano plots highlighted the pattern of fewer gene regulation differences between control cells and *E. rectale* CFSM, with H_2_O_2_ exposed cells consisted of nearly entirely significantly upregulated genes compared with *E. rectale* CFSM, with pro-inflammatory and host repair genes like FOS (72, 73), MUC15 (74), EGR1 (75), SLC38A3 (76, 77) (Figure 6B). GSH-treated cells displayed a different pattern with upregulation of some pro-inflammatory and repair genes, including C1S (78), PLAU (79), PPP1R15A (80), and downregulation of genes, including DAB2 (81), CKAP (82), and HHMR (83), which are involved in downregulation of the immune response, environmental stress sensing, and cell repair or migration, respectively. When comparing GSH treated cells with *E. rectale* CFSM treated cells, we observed several stress response genes upregulated in GSH treated cells similarly to *E. rectale* CFSM treated cells compared to inflamed control cells (Figure 6B). This result clearly indicates *E. rectale* CFSM alleviates host damage from H_2_O_2_, promotes host homeostasis and downregulates the immune response to cell damage.

To compare host expressed functions between treatment groups, gene counts between all treatment groups revealed upregulation of host damage and stress genes in damaged cells, including NECTIN (84), NGFR (85), MUC15, and EGR1. This immune response activation increases mucin production and activation of goblet cells. These genes were all expressed less in damaged cells treated with *E. rectale* CFSM, except for NECTIN1, but not downregulated in the H_2_O_2_ cells treated with 15 mM GSH (Figure 6C). However, both *E. rectale* CFSM and GSH treated cells displayed a differential pattern with upregulation of several stress and repair genes that were not upregulated in H_2_O_2_ exposed cells. Most genes upregulated in the *E. rectale* CFSM treatment are NECTIN1, NGFR, and ERC1, whereas GSH treated cells were lower expressed in those genes, indicating possible differences in immune activation between these treatments. GSH treated damaged cells were highly upregulated in genes involving environmental sensing and solute transfer, like ATF3, SPRR2G, PGF, and heat shock protein genes, while displaying downregulation of genes NECTIN1, ERC1, CDK6 and NGFR (Figure 6C).

Our work clearly shows that *E. rectale* microbial products were effective in reducing cell damage and repair gene classes. Our results not only quantify the effectiveness of GSH as a well documented antioxidant and promoter of cell homeostasis (86), but further demonstrate that *E. rectale* produces multiple bacterial metabolites that function synergistically as antioxidants to alleviate inflammation and, driving a balanced state, promoting gut repair. Although this pattern was unique from *E. rectale* CFSM, our work also showcases the genetic variations among individual microbial species, and its impact on their derived products, highlighting the implications of investigation of microbial populations and metabolites on a community level. We identify a product of *E. rectale*, reduced glutathione, as an antioxidant, as well as potential reigns on the immune response during ROS induced stress.

## Discussion

A deeper understanding of the gut microbiota and its impact on host inflammation and promotion of homeostasis is critical in developing treatments for colitis and associated inflammatory conditions. The Lachnospiraceae family has had a tumultuous history with nomenclature reclassifications; there has also been a fundamental lack of research seeking to understand pathways of individual microbial metabolites with the host and identification of microbial derived products that protect from, or alleviate excessive inflammation. In this study, longitudinal metagenomic sequencing of vertically transmitted dysbiotic mice revealed a potential role of the presence of Lachnospiraceae with recovery of induced dysbiosis and its resulting inflammation.

Treatment of colitis is challenging; dysbiosis is difficult to diagnose, and often requires patient-specific diagnostic and therapeutic regimens. With the safety concerns of fecal microbiota transplantation (FMT) (87, 88), an alternative approach could be using a community of commensal microbes to re-establish healthy host-microbe interactions and immune modulation. To assemble a community of microbes to facilitate gut health, a thorough understanding of individual microbes’ contributions or potential harm is necessary. Many Lachnospiraceae family members are known for their probiotic qualities in producing SCFA, reducing immune or inflammatory cell activation, and producing a variety of amino acids necessary for healing and recovery after damage in the gut. However, some species of Lachnospiraceae, including the genera *Blautia* and *Lachnospiraceae*, are associated with increased inflammation leading to disorders like ulcerative colitis, metabolic disorder, and chronic kidney disease (10). For the identification of microbes or microbial derived products for the treatment of chronic inflammatory conditions, a fundamental understanding of individual microbes and their products is essential.

Our results demonstrate *E. rectale* microbial products have the potential to alleviate ROS stress equivalent to that of 15 mM of GSH. GSH is a known ROS alleviator in rats (89) and is hypothesized as a suppressor of harmful inflammation in humans; however, dosage of up to 1000 mg (equivalent to 3.25 mM) a day has been shown to be ineffective in lowering oxidative stress markers (90). Highlighting the importance of microbial derived metabolites, we showed that *E. rectale* was also able to lower ROS levels during DSS induced colitis, while GSH alone was not sufficient in lowering ROS in inflamed cells. Our study addresses both at the level of potential gene function during glutathione biosynthesis, and at the metabolomic profile of *E. rectale*, revealing microbial products and pathways from *E. rectale* that may contribute to lowering inflammation and host damage. In addition, RNA sequencing of peroxide induced cells treated with *E. rectale* CFSM lowered environmental stress sensing genes and mucin production, indicating alleviation of ROS stress. While previous studies have explored the potential benefits of GSH (35, 91, 92), investigators focused on host-derived GSH or dietary supplementation, and have not adequately addressed microbial metabolites in lowering host experienced ROS stress. This study addressed the critical knowledge gap that identified glutathione as a microbial product of *E. rectale*, and its potential as a probiotic candidate for therapeutic intervention during gut inflammation.

## Materials & Methods

### STAR METHODS

● Experimental Models and Details

○ IL-10 KO Mice
○ Murine Serum
○ Murine Colon Histopathology
○ Murine Colon RNA sequencing
○ Bacterial Cultivation
○ Cell Culture
○ Lachnospiraceae Derived Metabolite Challenge to Inflamed Caco-2 Cells
○ Amino acid and Antioxidant Challenge to Inflamed and Damaged Caco-2 Cells
○ Metabolomics of Cell Supernatant
○ RNAseq of Caco-2 with Lachnospiraceae Derived Metabolites
● Bioinformatic workflow

○ Metagenomics
○ Human metagenomic analysis
○ BV-BRC analysis
○ Murine RNASeq analysis
○ Caco-2 RNASeq analysis
○ Statistical analyses
● Data Availability

○ Accession number and figshare

### Sample collection and processing

#### IL-10 KO Mice

All mice in this study were IL-10 KO Jax stock #002251/ C57BL/6J genetic background, consistent with other studies trying to emulate human gut inflammation in a mouse model (93, 94). Mice were bred in the Kansas State University Johnson Cancer Research Center mouse facility under *Helicobacter hepaticus*-free conditions, with a 12-hour light/dark cycle and up to four littermates of the same sex in a cage. We conducted this experiment using two different experimental sets of mice - Control mice (Control); mice without vertically transmitted dysbiosis, and mice with inherited dysbiosis (Dysbiotic), to investigate the impact of the resilient microbial populations on inherited dysbiosis. We supplemented dams with autoclaved drinking water with cefoperazone sodium salt (CPZ; 0.5 mg/mL), a broad spectrum antibiotic, when they were in the third week of gestation, to instill inherited dysbiosis (95). The duration of CPZ exposure lasted until pups were weaned at 21 days of age. Mice were monitored for signs of colitis daily. By week 23, remaining mice received 2.5% DSS for 5 days in autoclaved drinking water, and was changed every three days. We monitored mice daily during the time of DSS treatment and after treatment, until the end of the study. After a 16-day recovery period after 2.5% DSS treatment, all mice were sacrificed and tissues were collected.

#### Ethics Statement

All mouse experiments were reviewed and approved by the Institutional Animal Care and Use Committee at Kansas State University (APN #4391).

#### Murine Serum

We collected blood via heart puncture, and centrifuged to collect serum. Serum was stored in -80°C until further analysis. Mouse Cytokines were obtained using a 31-Plex assay (Eve Technologies, Canada, catalog: #33836) that measured Eotaxin, G-CSF, GM-CSF, IFN-γ, IL-1α, IL-1β, IL-2, IL-3, IL-4, IL-5, IL-6, IL-7, IL-9, IL-10, IL-12 (p40), IL-12 (p70), IL-13, IL-15, IL-17A, IP-10, KC, LIF, LIX, MCP-1, M-CSF, MIG, MIP-1α, MIP-1β, MIP-2, RANTES, TNF-α, and VEGF-A (not validated).

#### Murine Colon Histopathology

A section of the lower colon from each mouse was routinely fixed in 10% neutral-buffered formalin, trimmed, embedded in paraffin, sectioned, and stained with hematoxylin and eosin. Tissues were scored for characteristics of inflammation including surface epithelial damage, type and number of infiltrating inflammatory cells, and gland damage including dilation, inflammation, and hyperplasia. Each category was graded on a scale from 0 (normal), 1 (mild), 2 (moderate), or 3 (severe). Summative scores for each mouse and means for each treatment group were calculated.

#### Murine Colon RNA Sequencing

We stored a portion of the lower large intestine in TRIzol reagent (-80°C) until sequencing. Samples were sent to CD Genomics (Shirley, New York), where they were extracted and QC analysis performed by manufacturer protocols. Next, cDNA library preparation (poly-A selection) was completed for the 12 samples, and were then sequenced via NovaSeq PE150 for a total of 20 million pair-end reads.

#### Bacterial Cultivation

We obtained 5 strains of Lachnospiraceae members from ATCC: *Anaerostripes hadrus* strain B2-52, *Blautia coccoides* strain CLC-1, *Blautia producta* strain VPI 4299, *Eubacterium rectale* strain VPI 0990, and *Lachnoclostridium phytofermentans* strain ISDg. We cultivated these strains anaerobically (Coy Labs) in a gas mix (10% H2 10% CO2/N2) for 48 hours in chopped meat media using manufacturers’ suggestions (Anaerobic Systems).

#### Cell Culture

We obtained colorectal adenocarcinoma cells (Caco-2) cells from ATCC, and maintained them at 37°C with an atmosphere of 5% CO_2_, 95% Air, in 100 cm culture dishes. We maintained the Caco-2 cells in routine cell media consisting of filter sterilized Gibco Advanced DMEM supplemented with 10% heat inactivated Fetal Bovine Serum, 1% 10^5^ U/L penicillin, and 100 mg/L streptomycin, 1% Amphotericin B (fungizone). Cell generations 8-70 were used for this study with routine image comparisons ensuring no change in visible morphology. For experiments, we plated cells at 1 x 10^6^ cells per well, and grew to confluence with media changes 3 times per week. To induce inflammation, we introduced cells to 5% high molecular weight Dextran Sulfate Sodium (DSS) (MP Biomedicals) at 3 hours and 24 hours.

#### Lachnospiraceae Derived Metabolite Challenge to Inflamed Caco-2 Cells

We plated Caco-2 cells (generations 25-70) at 1 x 10^6^ cells per well in 24 well tissue culture plates, and grew to confluence up to 7 days with media changes 3 times per week. We collected bacterial spent media cultures and centrifuged them for 10 minutes at 10,000 x g, with supernatant collected, filter sterilized, and diluted in DMEM with DSS for all experiments. Cells were washed, with all media aspirated and 5% final DSS solution or 5 mM of hydrogen peroxide in DMEM and bacterial spent media were placed on the cells. We incubated the cells for 3 hours, and performed the Promega Griess Assay, measuring nitric oxide (as nitrite) according to the manufacturer’s protocol. We also performed the Promega LDH non-toxic assay according to the manufacturer’s protocol at 6 hour timepoints with both assays read on the Accuris smartreader 96. Finally, the DCFDA H2DCFDA Cellular ROS Assay Kit (ab113851) was used to measure intracellular ROS levels, with manufacturer’s protocols followed at 6 hours post introduction and measured using a Biotek Synergy H1 reader.

#### Metabolomics of Cell Supernatant

Caco-2 cells were seeded in 24-well plates grown to confluence for 7 days. On the day of the metabolite exposure, we incubated the cells in 1% DSS in cell culture media for 3 hours. We incubated with spent media from each Lachnospiraceae strain that were grown for 72 hours anaerobically in BHI broth, spun down and separated. We then added spent media, and cells were incubated for 3 hours under cell culture conditions. 0.5 mL of the cell culture supernatant was frozen at -80°C until analysis. Samples were extracted in 1:4 Methanol for LC-MS analysis (Metaware Bio, Woburrn, MA). Ultra Performance Liquid Chromatography (UPLC) and Quadrupole-Time of Flight Spectrometry were performed for widely targeted detection.

#### RNAseq of Caco-2 with Lachnospiraceae Derived Metabolites

We seeded the caco-2 cells at 0.3 x 10^6^ cells per well in 6-well plates overnight using generations 29-50. We then washed the plates with buffer, and treatments were added to the wells and incubated for 6 hours in cell culture conditions. All cells except the control received 5 mM of hydrogen peroxide, in addition to the treatment of *E. rectale* spent media, 15 mM GSH or chopped meat media that *E. rectale* was cultivated in. After the incubation period, we washed the cells with a cold buffer on ice, and scraped in the buffer. Cells were then spun down at 1600 rpm and the supernatant was removed. Pellets were then flash frozen and stored in -80°C with RNA extraction and sequencing performed by CD Genomics (Shirley, NY).

### Bioinformatic workflow

#### Metagenomics

We automated the metagenomics bioinformatics workflows using the program ‘anvi-run-workflow’ in anvi’o ver 7.1 (96). These workflows use Snakemake (97), and are equipped with numerous tasks including short-read quality filtering, assembly, gene calling, functional annotation, hidden Markov model search (98), metagenomic read-recruitment, and binning (99). We used the program ‘iu-filer-quality-minoche’ to form short metagenomic reads, and eliminate low-quality reads according to the criteria outlined by Minoche et al. (100). We used MEGAHIT v1.2.9 (101) to co-assemble quality-filtered short reads into longer contigs (contiguous sequences). We assigned the metagenomes into 2 experimental groups (control/no-CPZ, n= 26; CPZ/no-gavage, n= 8) for the co-assembly. We then used the following designs to process the assemblies: we used (1) ‘anvi-gen-contigs-database’ on contigs to compute k-mer frequencies and identify open reading frames (ORFs) using Prodigal v2.6.3 (102); (2) ‘anvi-run-hmms’ to identify sets of bacterial and archaeal single-copy core genes using HMMER v.3.2.1 (103); (3) ‘anvi-run-ncbi-cogs’ to annotate ORFs from NCBI’s Clusters of Orthologous Groups (COGs) (104); and (4) ‘anvi-run-kegg-kofams’ to annotate ORFs from KOfam HMM databases of KEGG orthologs (105). We mapped metagenomic short reads to contigs using Bowtie2 v2.3.5 (106) and converted to BAM files using samtools v1.9 (107). We profiled the BAM files using ‘anvi-profile’ with minimum contig length of 1000bp. We used ‘anvi-merge’ to unite all profiles into an anvi’o merged profile for all downstream analyses. We then utilized ‘anvi-cluster-contigs’ to group contigs into initial bins with CONCOCT v1.1.0 (108), and used ‘anvi-refine’ to manually curate the bins based on tetranucleotide frequency and contrasting coverage across the samples. We marked all bins that were > 70% complete and < 10% redundant as metagenome-assembled genomes (MAGs). Finally, we used ‘anvi-compute-genome-similarity’ to determine the average nucleotide identity (ANI) of our genomes using PyANI v0.2.9 (109), and identified non-redundant MAGs. We then conducted all downstream analyses on only non-redundant MAGs. We used the “detection” metric to assess occurrence of MAGs in a given sample. Detection of MAGs is the proportion of the nucleotides that were covered by at least one short read. We considered a MAG was detected in a metagenome if the detection value was > 0.25, to eliminate false-positive signals in read recruitment results, for its genome.

#### Human metagenomic analysis

We obtained the genomes from the discovery PRISM cohort (NCBI BioProject number PRJNA400072) with 161 adult patients enrolled with either ulcerative colitis (UC) or Crohn’s Disease (CD) and non-IBD control adults (49). We then mapped the five ATCC Lachnospiraceae genomes we obtained for molecular testing to the PRISM cohort genomes using Bowtie2 v2.3.5 (106) and samtools v1.9 (107) for converting to BAM files.

#### BV-BRC analysis

We uploaded and annotated all non-redundant MAGs in Bacterial and Viral Bioinformatics Resource Center (BV-BRC) (110, 111). All 73 MAGs resolving to the Lachnospiraceae family were grouped, and for each MAG, we used the Subsystem genes for comparison of potential functions and virulence factors between the Lachnospiraceae MAGs related to the ATCC cultures obtained. Counts of genes by each class of interest were used to build a heatmap and pathway constructs for each MAG of interest by using the Pathway function in BV-BRC linked to KEGG.

#### Murine RNASeq analysis

We used FastQC (112) and multiQC (113) to check the raw reads’ quality, and used Trimmomatic (114) to trim the reads. We used Trinity ver 2.13.1 (115) for de novo assembly, and RSEM (116) and DESeq2 (117) to estimate the expression levels and differential expression analysis respectively. Annotation was carried out using blast+ 2.11 and R version 4.0.0. The transcripts compared in DESeq2 were first blasted against mouse transcript with C57BL/6NJ as a reference ensemble ID (http://ftp.ensembl.org/pub/release-107/fasta/mus_musculus_c57bl6nj/cdna/), then the top hit (lowest e-value and largest length with minimum of 150bp) were kept. Finally, we used GO_MWU (118) to analyze the annotated transcripts.

#### Caco-2 RNAseq analysis

A nf-core/RNAseq pipeline v3.12.0 based on nextflow (v 23.04.3.5875) with a singularity container was chosen for the RNA data analysis. Default parameters and tools are used in the pipeline except that the raw reads were mapped with STAR and quantified using RSEM (116) with the reference genome GRCh38 release-110. DESeq2 (117) package was used for the downstream statistical analysis. Gene ontology (GO_MWU) enrichment analysis (118) was used to analyze annotated transcripts.

### Statistical analyses

Differences between groups were analyzed by one-way ANOVA with Tukey HSD test, unless stated otherwise. P-values of less than 0.05 were considered statistically significant. Statistical analyses were performed in RStudio. We used the program - Data Integration Analysis for Biomarker discovery using Latent cOmponents (DIABLO) to build a model to identify key components in the metagenomic data and host RNASeq data (119, 120). We (i) first identified key omics variables in the integration process; (ii) and then maximized the common and correlated information between the datasets; and (iii) finally visualized the results to identify relevant microbial and host genomic markers. Univariate and multivariate analyses were performed on metabolomics data to identify the metabolites from different treatments (Metaware Bio, Woburrn, MA).

### Data Availability

#### Accession Number and figshare

Raw sequencing data for the mice metagenomes were uploaded to SRA under NCBI BioProject PRJNA1074399. All other analyzed data in the form of databases and fasta files are accessible at figshare https://doi.org/10.6084/m9.figshare.25702764.v1.

## Acknowledgements

We appreciate Clark Bloomer and the entire team with the Genome Sequencing Facility at the University of Kansas Medical Center for help with all shotgun metagenomic sequencing in this study. We would like to acknowledge Dr. Tracy Meisner for services in technique and experimental design. Our thanks to Dr. Sherry Fleming with the Johnson Cancer Center at Kansas State University for support and helpful discussion. We also acknowledge the Johnson Cancer Center at Kansas State University for funding and support for this project as well as personnel. Finally, we would like to acknowledge the 34 IL-10 KO mice included in this study for their sacrifice to build our understanding of persistent gut inflammation and dysbiosis.

## Author Contributions

T.G.R. and S.T.M.L. designed this study. Sample collection was performed by T.G.R., C.I., L.H., A.K., S.P., K.M., T.S., H.W., S.S. Authors T.S., T.G.R., L.H., and H.W. completed DNA extraction including Nanodrop and Qubit quality analysis. Q.R. performed all RNAseq bioinformatic analysis. T.G.R. and S.T.M.L. performed anvi’o and BV-BRC bioinformatic analyses. Bacterial and cell culturing and isolation was done by T.G.R., H.W., and C.I.. A.S. and E.C. provided support for metabolite profiles. Authors T.G.R., H.W., C.I., and S.T.M.L. attributed biological relevance, wrote the manuscript, prepared figures, and supplementary files. T.G.R., S.T.M.L., and B.L.P. performed major manuscript and figure refinement while remaining authors contributed to lighter refinement. B.L.P. performed all histological analyses. S.T.M.L. acquired fundings for this study. All authors read, contributed to manuscript revision, and approved the submitted version of this manuscript.

## Supplemental Legends

Supplementary Table 1. Murine study information with weights by week as well as histology and serum cytokine results.

Supplementary Table 2. Shotgun metagenomic sequencing data including quality control, MAG summary and detections across murine samples.

Supplementary Table 3. Lachnospiraceae MAGs metabolic and gene comparison data as well as human patient mapping results.

Supplementary Table 4. Cellular health assay results including normalized results, and statistics, listed by stressor and assay type.

Supplementary Table 5. Metabolomics results for media control samples, and all experimental samples.

Supplementary Table 6. RNA sequencing results, counts data, results for heatmap, and DESeq2 results for each comparison used for volcano plots.

Supplementary Figure 1. Clustered heatmap displaying all detected, nonredundant MAGs across all murine samples throughout longitudinal study.

Supplementary Figure 2. Gene ontology (GO) analysis trees indicating differentially regulated gene classes by comparison. Red text is upregulated in the first comparison group, blue is downregulated in the first comparison group. Significance is marked by font size and italicized or bolded text.

