## Supplementary figures and images for "*Eubacterium rectale* detoxification mechanism increases resilience of the gut environment"

### Supplementary Figure S1

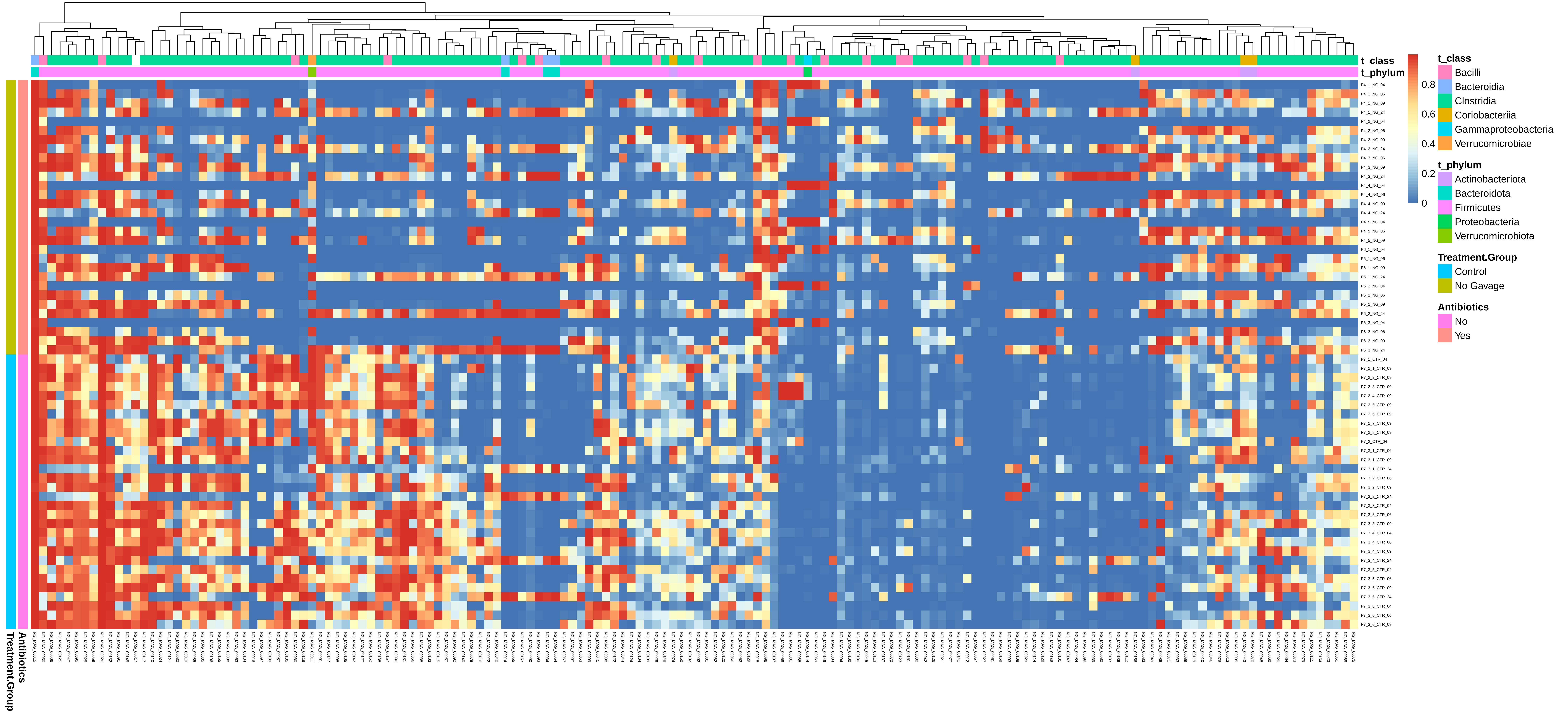

### Supplementary Figure S2

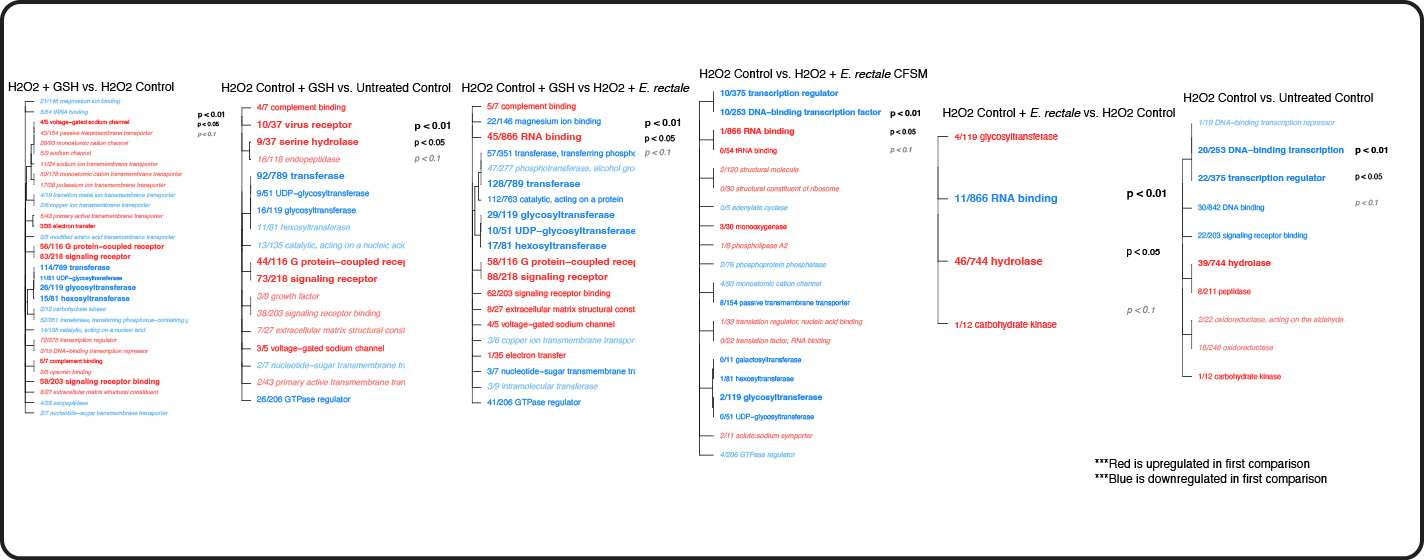
